# GSK3 activity is a cell fate switch that balances the ratio of vascular cell type

**DOI:** 10.1101/2020.02.08.939595

**Authors:** Takayuki Tamaki, Satoyo Oya, Makiko Naito, Yasuko Ozawa, Tomoyuki Furuya, Masato Saito, Mayuko Sato, Mayumi Wakazaki, Kiminori Toyooka, Hiroo Fukuda, Ykä Helariutta, Yuki Kondo

**Author notes:** **<u>Contact</u>**, Corresponding author: Yuki Kondo, Department of Biological Sciences, Graduate School of Science, The University of Tokyo, 7-3-1 Hongo, Bunkyo-ku, Tokyo, 113-0033.

## Abstract

The phloem transports photosynthetic assimilates and signalling molecules. It mainly consists of sieve elements (SEs), which act as “highways” for transport, and companion cells (CCs), which serve as “gates” to load/unload cargos. Though SEs and CCs function together, it remains unknown what determines the ratio of SE/CC in the phloem. In this study, we develop a novel culture system for CC differentiation named VISUAL-CC, which reconstitutes the SE-CC complex formation. Comparative expression analysis in VISUAL-CC reveals that SE and CC differentiation tends to show negative correlation, while total phloem differentiation is unchanged. This varying SE/CC ratio is largely dependent on GSK3 kinase activity. Indeed, *gsk3* hextuple mutants possess much more SEs and less CCs in planta. Conversely, *gsk3* gain-of-function mutants induced by phloem-specific promoter partially increased the CC ratio. Taken together, GSK3 activity appears to function as a cell fate switch in the phloem, thereby balancing the SE/CC ratio.

## Introduction

Multicellular organisms possess a variety of functional cells with a proper ratio for their life maintenance. In plants, the phloem tissue is composed of major two cell types; sieve elements (SEs) as conductive tubes and phloem companion cells (CCs) as helper cells for phloem transport. Phloem CCs function to support neighbouring SEs through connected plasmodesmata. Although they function together with each other to ensure phloem transport, it has long been a deep mystery how the ratio of SE/CC is strictly controlled in the phloem. Recent technical advances enabled to identify various regulators that control SE differentiation^*1*^. In contrast to SEs, understanding of the molecular mechanism underlying CC differentiation remains a long-standing challenge.

Vascular cell Induction culture System Using Arabidopsis Leaves (VISUAL) is a culture system that can artificially mimic plant vascular development^*2*^. In the VISUAL system, mesophyll cells are reprogrammed into vascular stem cells, and then differentiated into xylem vessel elements or phloem SEs within a couple of days. VISUAL enables the molecular genetic studies of vascular development, leading to the in-depth understanding of regulatory network especially for phloem SE differentiation. Even in VISUAL, differentiation into phloem CCs rarely occurs^*2*^, which makes it difficult to study CC development in detail.

In this study, we develop a new culture system for CC differentiation named VISUAL-CC by modifying the conventional VISUAL method. Based on comprehensive gene expression analysis in VISUAL-CC, here we reveal that GLYCOGEN SYNTHASE KINASE 3 (GSK3) activity plays an important role in determining the SE/CC ratio. *In vivo* genetic analyses confirm the importance of GSK3 as a cell fate switch in phloem development.

## Main text

### VISUAL-CC is a novel culture system for CC differentiation

The conventional VISUAL system can induce ectopic xylem (XY) or phloem SEs via the stage of vascular stem cell (Fig. 1a). Toward the understating of CC differetniation, we modified the VISUAL based on a luciferase-based screen with *SUCROSE-PROTON SYMPORTER 2* (*SUC2*) *pro:ELUC*, a specific CC marker^*3*^ (Fig. 1b). In this screen process, vascular stem cells were induced by the conventional VISUAL method in advance, subsequently were exposed to a variety of culture media. After a series of screens with different media, we could induce *pSUC2:ELUC* activity (Fig. 1c) and ectopic expression of the *pSUC2:YFPnls* marker in cotyledons within 4 days using CC medium (Fig. 1d). Hereafter, we refer to this culture system as “VISUAL-CC”. To further investigate the spatial pattern, a dual phloem marker line expressing *pSUC2:YFPnls* and *SIEVE-ELEMENT-OCCLUSION-RELATED 1* (*SEOR1*) *pro:SEOR1-RFP^5^*, a specific SE marker was established. Detailed observation of the dual phloem marker line by confocal microscopy after tissue-clearing treatment (ClearSee)^*4*^ revealed that CCs expressing *pSUC2:YFPnls* (green) are detected only in dividing cell-clusters and are always limited to the cells adjacent to SEs expressing *pSEOR1:SEOR1-RFP^5^* (red) (Fig. 1e). Thus, CC and SE markers appeared next to each other after several rounds of cell division (Fig. 1e). Observations using a field emission scanning electron microscope (FE-SEM) or transmission electron microscopy (TEM) consistently visualized CC-like cells with dense cytoplasm adjacent to SEs with brighter cytoplasm (Fig. 1f). These cells showed minor vacuolation and developed the branched plasmodesmata typically seen in SE-CC complexes *in vivo* (Fig. 1f–h). In VISUAL, SMXL4 and SMXL5 are known as important regulators for early phloem SE development (Supplementary Fig. 1a)^*6, 7*^. In VISUAL-CC, the double mutant *smxl4 smxl5* significantly suppressed CC differentiation (Supplementary Fig. 1b), suggesting that SE and CC differentiation shares a common developmental process from vascular stem cells. Taken together, these results suggest that VISUAL-CC can reconstitute the SE-CC complex formation.

**Figure 1.**
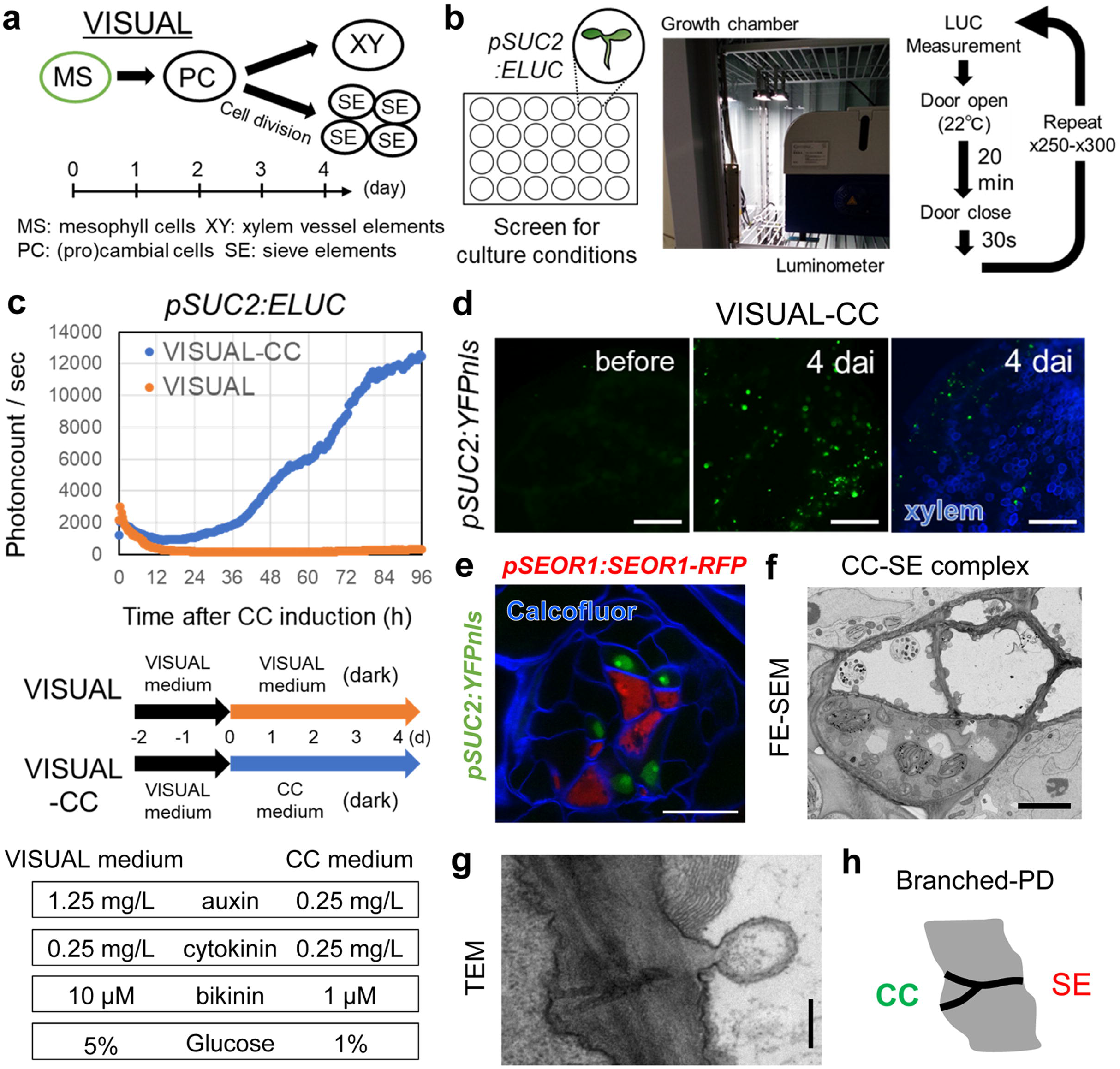
VISUAL-CC is a novel method for inducing SE-CC complexes. **a**, The process of vascular cell differentiation in conventional VISUAL. **b**, Schematic of the screening system. LUC activity of *pSUC2:ELUC* was monitored every 20 minutes during culture under various conditions. **c**, Time-course of *pSUC2:ELUC* intensities during culture in conventional VISUAL and VISUAL-CC. The vertical axis indicates photon counts per second detected by the luminometer. The lower panel illustrates culture conditions and medium composition during VISUAL and VISUAL-CC culture; see Materials and Methods for a detailed protocol. **d**, *pSUC2:YFPnls* expression before and after VISUAL-CC treatment. Xylem cells were detected using UV autofluorescence (blue). **e**, Expression patterns of *pSEOR1:SEOR1-RFP* (red) and *pSUC2:YFPnls* (green) in VISUAL-CC. Cell walls were stained with calcofluor white (blue). **f**, FE-SEM image of ectopic CC-like cells (high electron density) and SEs (low electron density) induced by VISUAL-CC. **g**, TEM image of branched plasmodesmata between an ectopic CC-like cell and a SE. **h**, Schematic of branched plasmodesmata. Scale bars: 500 μm (d); 5 μm (f); 200 nm (g)

### VISUAL-CC can induce known CC-related genes expression

To validate the promoter-based assay, we compared *SUC2* mRNA accumulation with the promoter:LUC activity in the same sample (Fig. 2a, b). qRT-PCR analyses of VISUAL-CC samples and samples cultured in the conventional VISUAL medium (VISUAL, V) as negative controls revealed a strong correlation between promoter activity and mRNA level of *SUC2* (Fig. 2b; r = 0.97, *P* < 0.005) (Fig. 2b). We used these data for classification of VISUAL-CC samples into strong LUC activity (CC-strong, S) and moderate LUC activity (CC-moderate, M), according to their *SUC2* levels (Fig. 2a, b; Supplementary Fig. 2). Expression of *SISTER OF ALTERED PHLOEM DEVELOPMENT* (*SAPL*), another CC marker gene^*8*^, also showed a strong correlation with *SUC2* expression (Fig. 2c; r = 0.91, *P* < 0.005). A microarray analysis using the same samples was performed to obtain a comprehensive gene expression profile (Supplementary Fig. 3a). As expected, genes previously characterized as CC-related, including *SULFATE TRANSPORTER 2;1* (*SULTR2;1*)^*9*^, *SODIUM POTASSIUM ROOT DEFECTIVE 1* (*NaKR1*)^*10*^, *C-TERMINALLY ENCODED PEPTIDE RECEPTOR* (*CEPR1*)*/ XYLEM INTERMIXED WITH PHLOEM 1* (*XIP1*)^*11, 12*^ and *MYB-RELATED PROTEIN 1* (*MYR1*)^*13*^, showed similar expression patterns to *SUC2* and *SAPL* (Fig. 2d). Consistently, quantitative RT-PCR assay for these genes validated the microarray result (Supplementary Fig. 4). By utilizing the variation in expression observed between samples (S, M, V), we identified 186 VISUAL-CC inducible genes that satisfied the following patterns of expression levels: S > M > V and S/V > 4 (Fig. 2E and Supplementary Fig. 2). According to the previous dataset of root cell type-specific transcriptome^*14*^, these genes were mainly expressed in root CCs or phloem pole pericycles (PPPs; Fig. 2f). PPPs are also known to participate in phloem unloading from SEs in roots via intervening plasmodesmata^*15*^ (Fig. 2f). Here we grouped 67 genes as VISUAL-CC-related (VC) genes based on CC-preferential expression (Supplementary Table 1). Transporter genes were over-represented among these VC genes, reflecting the functional aspect of phloem transport (Supplementary Fig. 3b, c).

**Figure 2.**
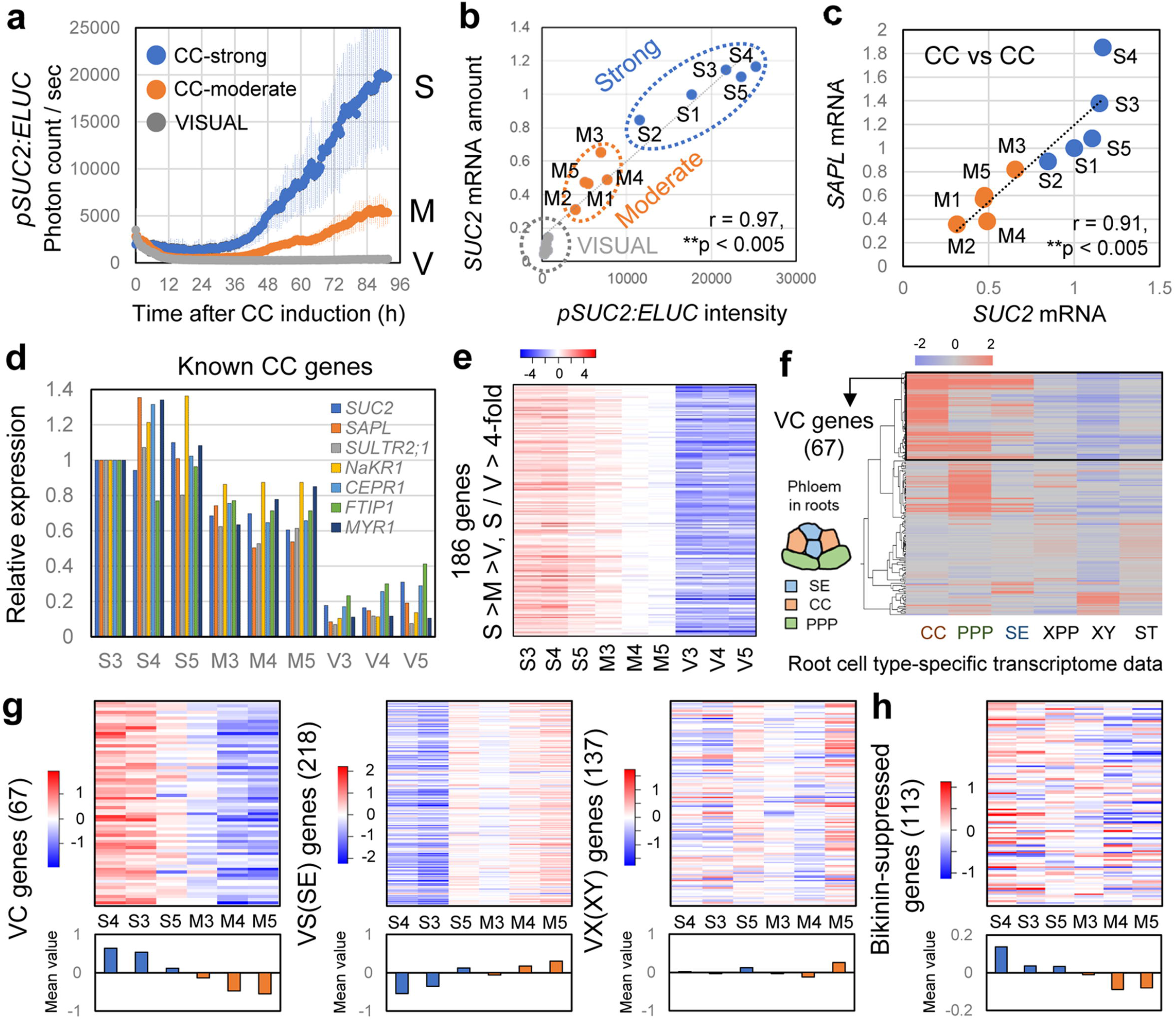
Transcriptome analysis in VISUAL-CC. **a**, Time-course of *pSUC2:ELUC* signal intensities for each category: CC-strong (S), CC-moderate (M), and VISUAL (V) (n = 5). Error bars indicate SD. **b**, Correlation between *pSUC2:ELUC* signals and relative *SUC2* mRNA expression levels for each sample. **c**, Correlation of expression levels between *SUC2* and *SAPL* in S and M samples. **d**, Expression levels of known CC-related genes in each sample. Relative expression levels were calculated when expression in S3 was set to 1. **e**, Heat map of expression levels of VISUAL-CC inducible genes in each sample. Expression data were normalized against the median value and are presented according to the color scale (log2) above the chart. **f**, Cluster analysis of VISUAL-CC inducible genes based on a previous root cell type-specific transcriptome dataset^14^. A schematic of the primary phloem tissue patterning found in roots is shown on the left-hand side of the chart. **g**, Heat map of expression levels of VC (67), VS (218), and VS (137) genes in S and M samples. The lower panel indicates the mean value for each sample. **h**, Heat map of expression levels of 113 bikinin-suppressed genes in S and M samples.

### SE and CC differentiation tends to show negative correlation in VISUAL-CC

We previously identified 137 VISUAL-XY-related (VX) genes and 218 VISUAL-SE-related (VS) genes using VISUAL microarray data^*2*^. Then, expression of VX and VS genes was examined in the VISUAL-CC transcriptome dataset. Although there was no regular pattern of VX gene expression, expression of VS genes was very low in the S samples in contrast to that of VC genes (Fig. 2g). Correlation analysis among these gene sets revealed that expression levels of VC genes negatively correlate with those of VS genes (Supplementary Fig. 5b; r = −0.91, *P* < 0.05) but not with those of VX genes (Supplementary Fig. 5a; r = −0.28, *P* > 0.05). To further assess this tendency, we calculated the quantitative expression levels of vascular marker genes in individual samples. Although there was a strong correlation between expression of *SAPL* (CC) and *SUC2* (CC) (Fig. 2c; r = 0.91, *P* < 0.005), no correlation was found between *IRREGULAR XYLEM 3* (*IRX3*)^*16*^ (XY) and *SUC2* (CC) expression (Fig. 3a; r = −0.27, *P* > 0.05). By contrast, expression of *SEOR1* (SE) showed a significant negative correlation with that of *SUC2* (CC) (Fig. 3b; r = −0.76, *P* < 0.05). All these results suggest that CC differentiation and SE differentiation tend to show negative correlation. Interestingly, expression levels of *ALTERED PHLOEM DEVELOPMENT* (*APL*) (SE+CC), which is expressed in both SEs and CCs^*17*^ was almost constant among all the samples, indicating that total amount of differentiating phloem cells is almost unchanged (Fig. 3c and Supplementary Fig. 6a). Taken together, these results suggest that VISUAL-CC induces different ratios of CC-like cells and SEs without changing the total number of phloem cells, while VISUAL only produced SEs. This implies that a key determinant of SE or CC cell fate is present in VISUAL-CC cultures.

**Figure 3.**
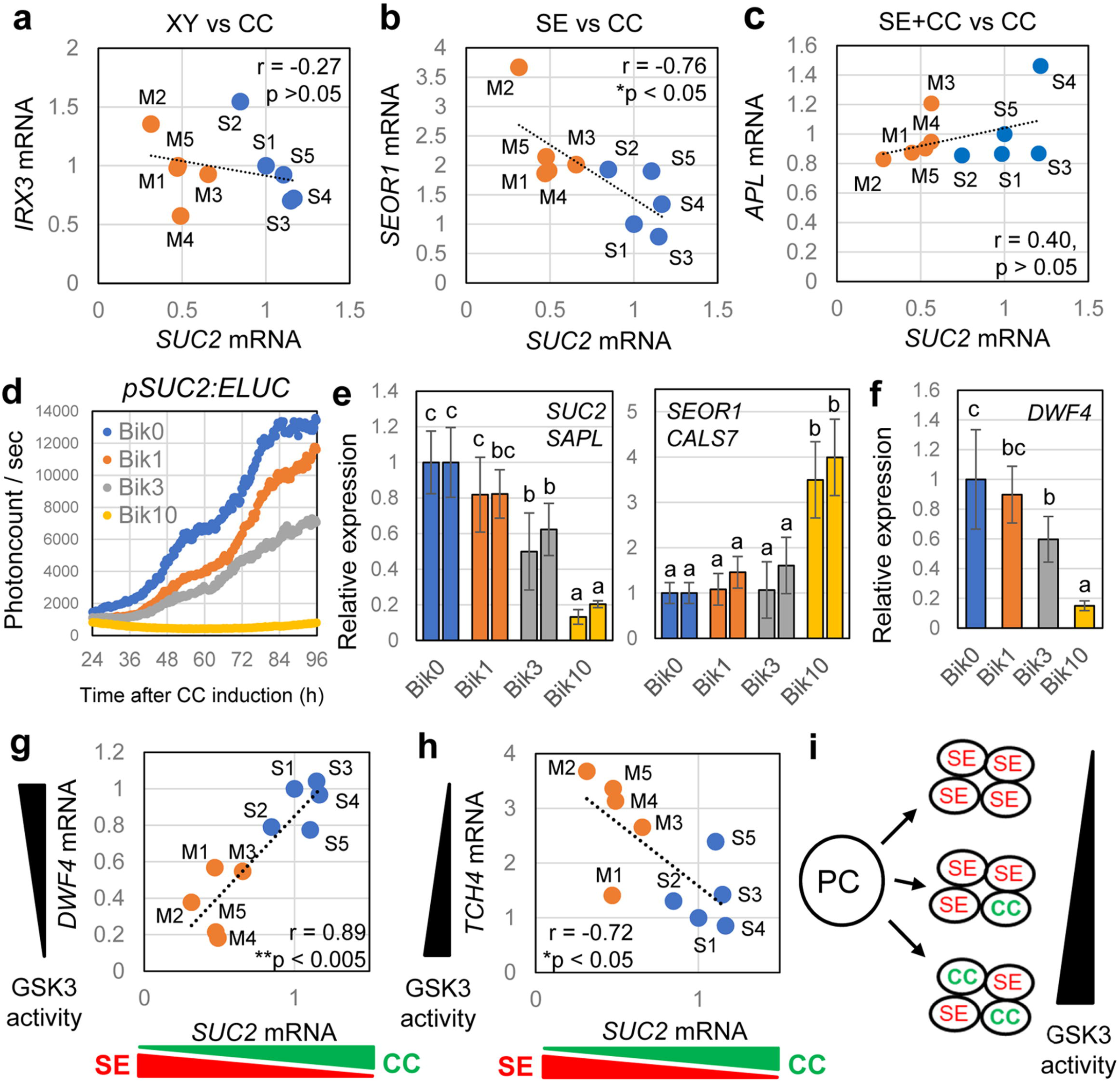
GSK3s activity balances the SE/CC ratio in VISUAL-CC. **a**, Expression of *SUC2* (CC) and *IRX3* (XY) showing no correlation between levels in S and M samples. The Pearson correlation coefficient and *P*-value are marked on the chart. **b**, Negative correlation between expression of *SUC2* (CC) and *SEOR1* (SE). **c**, Positive correlation between expression of *SUC2* (CC) and *APL* (SE+CC). **d**, Time-course of *pSUC2:ELUC* signal intensities during VISUAL-CC with various bikinin concentrations. Averaged LUC values are shown (n = 6). **e** and **f**, Expression of *SUC2*, *SAPL*, *SEOR1*, *CALS7*, and *DWF4* in VISUAL-CC samples cultured with various bikinin concentrations for 4 days. Statistical differences between samples are indicated by different letters (ANOVA, Tukey-Kramer method; n = 6; error bars indicate SD). **g**, Positive correlation between expression of *SUC2* (CC) and *DWF4* (GSK3-induced). **h**, Negative correlation between expression of *SUC2* (CC) and *TCH4* (GSK3-supprressed). **i**, Schematic model showing dose-dependent regulation of the SE/CC ratio by GSK3 activity.

### The SE/CC fates are determined mostly depending on GSK3 activity

To identify the determinants of SE/CC cell fate, we investigated the impact of auxin and bikinin, as their concentrations differed between VISUAL and VISUAL-CC media (Fig. 1c). Exogenous auxin reduced *pSUC2:ELUC* expression in a dose-dependent manner (Supplementary Fig. 7a). Although *SUC2* expression levels showed a similar decreased trend (Supplementary Fig. 7b), *SEOR1* expression was not affected by auxin level (Supplementary Fig. 7c). Similarly, bikinin showed a repressive effect on *pSUC2:ELUC* expression levels (Fig. 3d). Unlike auxin, treatment with bikinin decreased expression of *SUC2* in a dose-dependent manner while simultaneously increasing *SEOR1* expression (Fig. 3e). Furthermore, two other CC and SE markers, *SAPL* and *CALLOSE SYNTHASE 7* (*CALS7*)^*18*^ showed a similar response against bikinin treatment (Fig. 3e). Bikinin is a specific inhibitor of plant GSK3s^*19*^ and thus these results suggest that GSK3 activity plays a role in determining the SE/CC ratio. GSK3 activity correlates with the expression of brassinosteroid biosynthetic genes, as a negative feedback regulation. Indeed, *DWARF4* (*DWF4*)^*20*^, one of brassinosteroid biosynthetic genes, was down-regulated following bikinin treatment in a dose-dependent manner (Fig. 3f). In the VISUAL-CC transcriptome, expression of brassinosteroid biosynthetic genes tended to be higher in the S samples and lower in the M samples (Supplementary Fig. 8), also suggesting the relationship between the GSK3 activity and the SE/CC ratio. Previous studies have reported 113 bikinin-suppressed genes^*19*^, then we investigated the correlation between these genes and VC genes in VISUAL-CC transcriptome data. The bikinin-suppressed genes showed higher expression in the S samples (Fig. 2h) and their expression exhibited positive correlation with expression of VC genes (Supplementary Fig. 5c; r = 0.94, *P* < 0.01). To investigate the relationship further, we quantitatively compared the expression patterns of *SUC2* and GSK3-affected genes in the S and M samples. Expression of *DWF4*, a typical GSK3-induced gene (Fig. 3f), showed a strong positive correlation with *SUC2* expression (Fig. 3g; r = 0.89, *P* < 0.005). Similarly, other GSK3s-induced genes such as *CONSTITUTIVE PHOTOMORPHOGENIC DWARF* (*CPD*) and *BRASSINOSTEROID-6-OXIDASE 2* (*BR6ox2*) showed significantly higher expression in the S samples than in the M samples^*21, 22*^ (Supplementary Fig. 6b). By contrast, expression of *TOUCH 4* (*TCH4*), a typical GSK3-supprressed gene^*23*^, showed a significant negative correlation with *SUC2* expression (Fig. 3h; r = −0.72, *P* < 0.05). These results strongly suggest that the SE/CC ratio is largely dependent on GSK3 activity *in vitro* (Fig. 3i).

### Genetic manipulation of GSK3 activity alters the *in vivo* SE/CC ratio

Then, we analysed the role of GSK3s in *in vivo* secondary phloem development in *Arabidopsis* hypocotyls. In hypocotyls, SEs are characterized by vacant cytoplasm whereas CCs are deeply stained with toluidine blue and they usually appear as pairs in a transverse section (Fig. 4a). Inhibition of GSK3 activity by bikinin treatment induced clusters of SEs and far fewer CCs (Fig. 4a). Bikinin treatment consistently reduced expression of *pSUC2:YFPnls* and resulted in clusters of *pSEOR1:SEOR1-RFP* signals in the dual phloem marker line (Fig. 4b, c), indicating that bikinin promotes SE formation and decreased CC number *in vivo*. Next, we confirmed the function of GSK3 proteins genetically using knock-out mutants of members of the SKII subfamily (*BIN2*, *BIL1*, and *BIL2*) and RNAi knock-down for SKI subfamily members (*AtSK11*, *AtSK12*, and *AtSK13*)^*24*^, because they are the main targets of bikinin^*17*^. The phloem tissue of the *bin2 bil1 bil2 AtSK13RNAi* quadruple mutant exhibited a slight but significant decrease in CC occupancy (40%) when compared with wild-type plants (44%) (Fig. 4d, e and Supplementary Fig. 9). The *gsk* hextuple mutant (*quadruple + AtSK11*, *AtSK12RNAi*) showed a reduction in CC occupancy (20%), resulting in more SEs and fewer CCs (Fig. 4d, e and Supplementary Fig. 9). Moreover, in the hextuple mutant, some of the PPP cells unexpectedly differentiated into ectopic SE-like cells (Fig. 4d, e). Previous studies have revealed that the vascular cells express SKII subgroup genes *BIN2* and *BIN2-LIKE2 (BIL2)^24^*. In addition, expression of SKI subgroup genes *pSK11:GUS* and *pSK12:GUS^25^* was found in the vasculature including the phloem tissue (Fig. 4f). Similarly to the GUS expression analysis, SKI/II genes expression was kept high in VISUAL time-course and in VISUAL CC transcriptome data (Supplementary Fig. 10), indicating that 6 GSK3 members are present during phloem development. Next we investigated local GSK3 activity in the vasculature using *pDWF4:GUS,* which is an indicator of high GSK3 activity. Supporting with our idea, *pDWF4:GUS* expression was detected in the phloem CCs but not in SEs (Fig. 4f). Taken together, our results indicate that GSK3 activity is required for maintaining high CC occupancy *in planta*.

**Figure 4.**
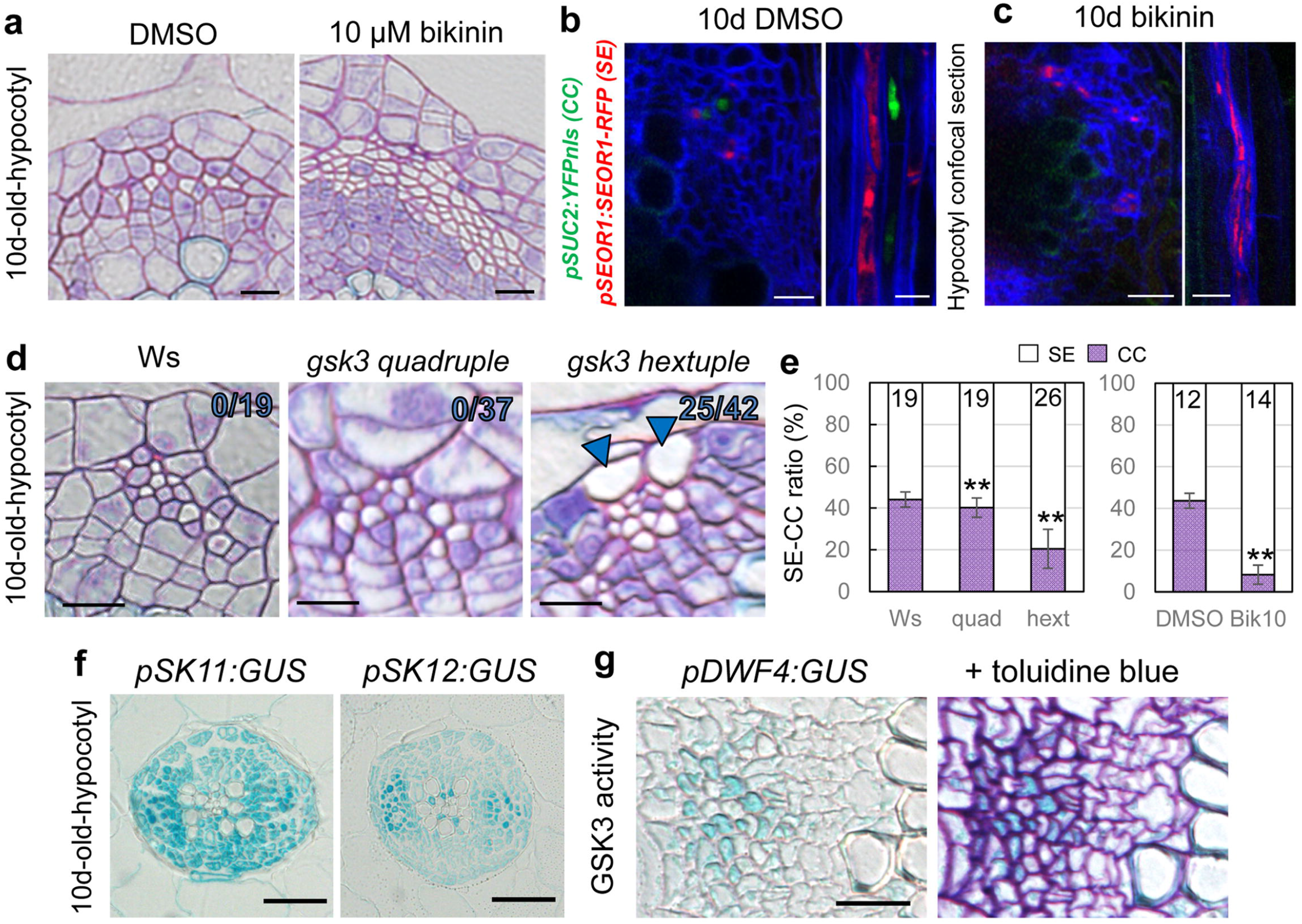
Reduction of GSK3 activity decreases the CC ratio *in planta*. **a**, Toluidine blue-stained transverse sections through mock-treated (DMSO) and bikinin-treated hypocotyls. SE: white empty cell; CC: dense purple cell. **b** and **c**, Expression of *pSEOR1: SEOR1-RFP* (red) and *pSUC2:YFPnls* (green) in 10-day-old hypocotyls treated with DMSO (b) or 10 μM bikinin (c) for 10 days (n = 8). Confocal images of transverse (left) and longitudinal sections (right). **d**, Toluidine blue-stained transverse sections for 10-day-old hypocotyls of Ws (wild-type) and *gsk3* high-order mutant plants. Arrowheads: ectopic SEs at the PPP position; the ratio of individuals showing ectopic SEs is marked on the upper right of the image. **e**, SE/CC ratios (%) in Ws, *gsk3* high-order mutants, and bikinin-treated plants calculated from toluidine blue-stained sections. Numbers of individuals are marked (n = 12–26). **f**, GUS staining of 10-day-old hypocotyls of *pSK11:GUS* and *pSK12:GUS* plants. **g**, GUS staining of 11-day-old hypocotyls of *pDWF4:GUS* plants. Left panel: GUS staining pattern on the cross section, Right panel: highly overlapping pattern of GUS and toluidine blue staining. Asterisks: significant differences determined using Dunnett’s or Student’s t-test (\*\**P* < 0.005). Scale bars: 10 μm (a–d); 20 μm (g); 50 μm (f)

GSK3s function as signalling hubs to control xylem differentiation in the cambium^*24*^. Here to focus on phloem development, *bin2-1*, a stable form of GSK3^*26*^, was driven under promoters specific to each stage of phloem development. As we had previously demonstrated that the sequential genetic cascade in phloem SE differentiation is *NAC020* (early), *APL* (middle), *SEOR1* (late)^*2*^, we induced expression of *bin2-1* under these different phloem promoters and investigated their phloem phenotype (Fig. 5a, b). Expression of *bin2-1* driven by the *APL* and *SEOR1* promoters did not affect phloem phenotypes, but *pNAC020:bin2-1* slightly but significantly increased the ratio of CCs in the phloem (Fig. 5a–c and Supplementary Fig. 9). To confirm the results with the CC marker, number of *pSUC2:YFPnls* signal in WT and *pNAC020:bin2-1* was quantified using confocal microscope (Fig. 5d). YFP-positive cell number estimated from 3D-reconstruction images was significantly higher in the *pNAC020:bin2-1* than in the WT (Fig. 5e, f). All these results indicate that GSK3s function as cell fate switches for determining differentiation into phloem CCs or SEs, and that GSK3 activity, especially in the early phloem development, was important for ensuring the proper ratio between CCs and SEs (Fig. 5g).

**Figure 5.**
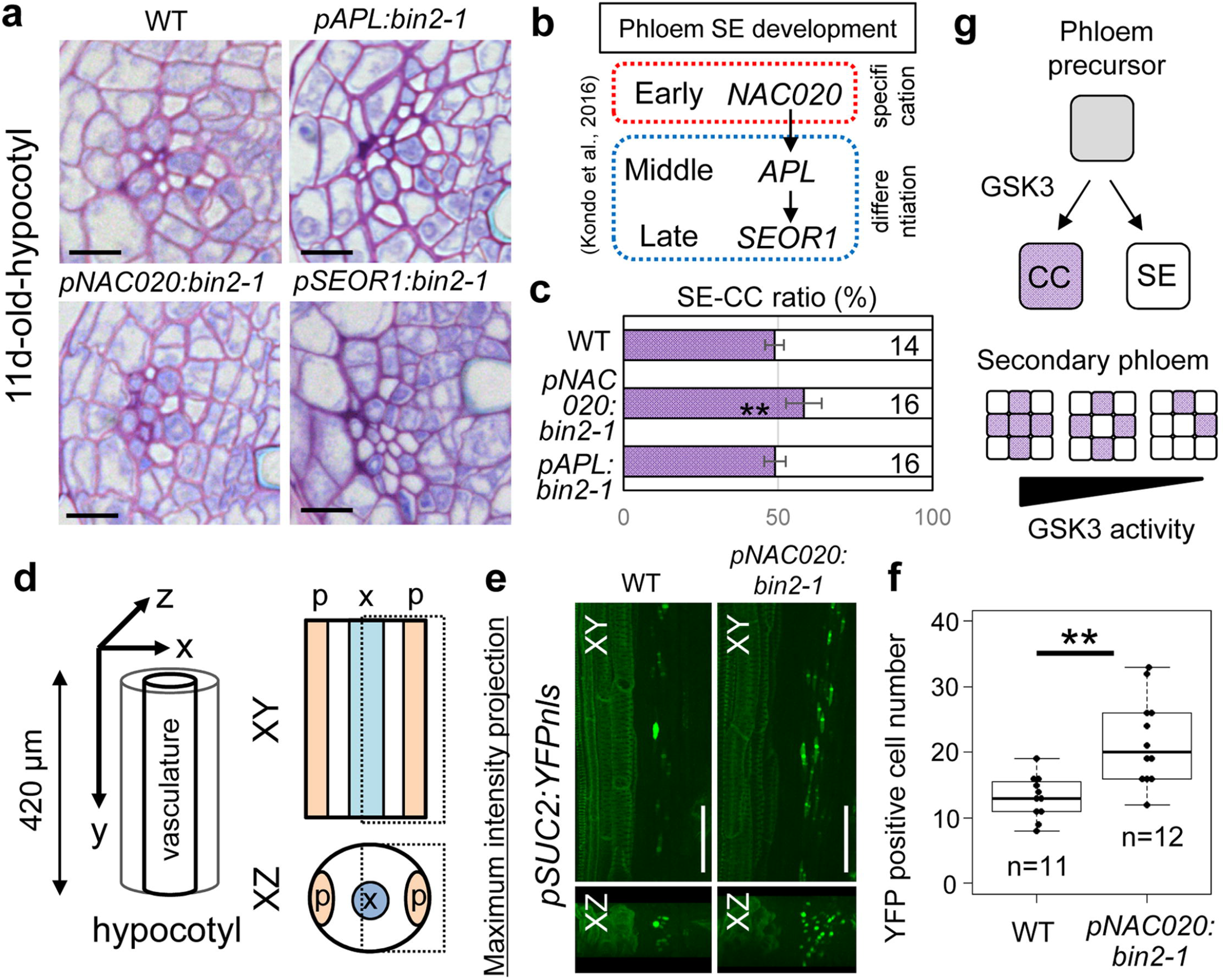
Early phloem-specific activation of GSK3 activity increases the CC ratio *in planta*. **a**, Toluidine blue-stained transverse sections through hypocotyls of 11-day-old Col (wild-type), *pNAC020:bin2-1*, *pAPL:bin2-1*, and *pSEOR1:bin2-1* plants. **b**, Genetic cascade revealed by VISUAL (n = 14–16). **c**, SE/CC ratios (%) in wild-type (WT), *pNAC020:bin2-1*, *pAPL:bin2-1* and *pSEOR1:bin2-1* calculated from toluidine blue-stained sections. **d**, Schematic illustration for 3D confocal imaging of *pSUC2:YFPnls* marker expression in hypocotyls. **e**, Maximum intensity projection of *pSUC2:YFPnls* marker expression in the WT and *pNAC020:bin2-1* was shown from XY (upper) and XZ (lower) angles. **f**, Quantification of *pSUC2:YFPnls* positive cells from 3D-reconstruction images in in the WT and *pNAC020:bin2-1* (n = 11–12). Asterisks: significant differences determined using Student’s t-test (\*\**P* < 0.005). **g**, Schematic showing that GSK3 activity determines SE/CC ratio. Scale bars: 10 μm (a); 50 μm (e)

### BR-BES1 signalling does not participate in SE/CC fate determination

Finally, we examined the involvement of brassinosteroid (BR) in CC differentiation, because GSK3s function as signal mediators in BR signalling^*26, 27*^. However, application of brassinolide (BL), an active BR, did not alter the SE/CC ratio in hypocotyls (Supplementary Fig. 11). Moreover, *bes1 bzr1* loss-of-function and *bes1-D bzr1-D* gain-of-function mutants for *BRI1-EMS-SUPPRESSOR 1* (*BES1*) and *BRASSINAXOLE RESISTANT 1* (*BZR1*), which are well-known transcription factors phosphorylated by GSK3s in BR signalling^*28*^, exhibited a normal phloem development in terms of the SE/CC ratio (Supplementary Fig. 12). These results suggest the possibility that other signalling pathway(s) than BR participates in controlling the SE/CC ratio.

## Discussion

Recent studies have revealed that CCs are not only of importance for phloem transport, but also act as a signal centre integrating environmental information into the developmental program^*29*^. We have established VISUAL-CC as a powerful tool for analysing the functional and developmental processes of CCs. During secondary vascular development, it has been widely believed that SEs and CCs are derived from the same phloem precursors via asymmetric cell division^*30*^. In VISUAL-CC, SEs and CCs were formed from vascular stem cells as neighbouring complexes after several rounds of cell division, suggesting that VISUAL-CC may reflect the process of secondary phloem development. By taking advantages of VISUAL-CC, we identified GSK3s as key molecular switches to specify cell fate toward SE or CC. Indeed, *gsk3* gain-of-function and loss-of-function mutants altered the ratio of SE/CC differentiation in the hypocotyl vasculature, and our expression analysis together with previous results consistently revealed that SKI/II subgroup members of GSK3s including BIN2 are expressed in the vascular tissues of hypocotyls^*24*^. As SEs lose their nuclei during the differentiation process, support from adjacent CCs is essential for their function. Thus, maintenance of the SE/CC ratio by GSK3s will be an important mechanism ensuring survival under various environmental conditions. Although GSK3s act as a central regulator of SE/CC development, genetic experiments suggested the involvement of other signalling than BR mediated by GSK3s in this regulation. GSK3s has been implicated in regulation of phloem development through the interaction with OCTOPUS (OPS), which genetically functions together with BREVIS RADIX (BRX)^*31*^. Further studies on such interacting proteins and reverse genetic approaches combined with VISUAL-CC transcriptome data will be helpful for elucidating a novel signalling cascade controlling the SE/CC ratio.

GSK3s also function in animals as molecular switches determining differentiation into alternate cell types^*32*^, suggesting their common and important role as cell fate switches. On the other hand, GSK3 activity regulates asymmetric cell division in the stomata lineage by interacting with polarly localized proteins, which are required for specifying stomatal cell fate^*33*^. Further investigation with the context of asymmetric cell division will uncover the extent to which GSK3s serve as a common mechanism determining cell fate.

## Methods

### Plant materials

Arabidopsis plants used in this study is Col-0 accession, except for *gsk3* high-order mutants (Ws background). To construct the CC-reporter lines, approximately 2.0 kb of the *SUC2* promoter region was cloned and then fused with ELUC (Toyobo) or YFP containing a nuclear localization signal. A genomic fragment of *SEOR1*, approximately 4.8 kb long and containing 1.6 kb of the promoter, was fused with RFP to make *pSEOR1:SEOR1-RFP*; this was subsequently transformed into *pSUC2:YFPnls* to generate the double phloem marker line. To construct *pDWF4:GUS*, approximately 1.9 kb of the *DWF4* promoter region was cloned and introduced into pMDC163 vector to create a *GUS*-fusion gene.

Phloem-specific GSK3 activation lines were constructed by cloning *bin2-1* (*BIN2E263K*) and fusing it with the *NAC020* (2.4 kb), *APL* (2.9 kb), and *SEOR1* (1.6 kb) promoters using the LR reaction (Thermo Fisher Scientific). The *gsk quadruple* and *hextuple* mutants (Ws accession) used in this study were as reported previously^*23*^. The *smxl4 smxl5* mutants were as reported previously^*6,7*^. The *bes1 bzr1* loss-of-function mutants and *bes1-D bzr1-D* gain-of-function mutants were as reported previously^*33*^. *pSK11:GUS* and *pSK12:GUS* lines were as reported previously^*25*^.

### LUCIFERASE measurement

In this study, ELUC with PEST domain (Toyobo) was used as a short half-life luminescent protein. *pSUC2:ELUC* seedlings were co-cultured with 200 μM D-luciferin (Wako) in white 24-well plates (PerkinElmer). The time-course of luciferase (LUC) activity was measured automatically using a TriStar2 LB942 (Berthold) within a growth chamber (Nihonika).

### Microscopic observation

For deep imaging with confocal microscopes, isolated tissue samples were fixed for 3 hours under vacuum in a fixative solution (4% paraformaldehyde and 0.01% Triton X-100 in 1× PBS). Fixed samples were washed twice with 1× PBS and transferred to ClearSee solution (25% urea, 15% xylitol, 10% sodium deoxycholate). ClearSee solution was replaced with fresh solution every 2 days for 3 to 4 weeks. Calcofluor staining was performed 1 week before microscopic observations by adding 0.1% (w/v) calcofluor white to the ClearSee solution. The samples were stained overnight and then washed with ClearSee solution without calcofluor. Once the samples were stained, washing was continued as described above. Cleared samples were observed using LSM880 (Zeiss) or FV1200 (Olympus) confocal microscopes with Z-stack. For the quantification of YFP-positive cells, we counted the number of cells in a phloem pole of approximately 420 μm length of hypocotyls based on reconstructed 3D images.

### Electron microscopy

Sample preparation for electron microscopy observation was modified slightly from a previous study^*35*^. Briefly, leaf disks induced by VISUAL-CC were fixed and embedded in resin. Thin sections (100 nm) were mounted on glass slides. Sections were stained with 0.4% uranyl acetate solution (UA) and a lead citrate solution (Pb), and then coated with osmium tetroxide. Observations of slides were made using a field emission scanning electron microscope (FE-SEM) (Hitachi SU 8220). Thinner (80 nm) sections were mounted on formvar-coated 1-slot copper grids, stained with 4% UA and Pb, and then observed using an 80 kV transmission electron microscope (JEOL JEM-1400 Flash).

### qRT-PCR and Microarray experiments

Total RNA was extracted from four cotyledons using RNeasy plant mini kit (Qiagen) after LUC measurement. After reverse transcription reaction, qRT-PCR was performed using LightCycler 480II (Roche) by a universal probe method. Expression value was normalized with an internal control *UBQ14*. Microarray experiments were conducted with the Arabidopsis Gene 1.0 ST Array (Affymetrix) and analyzed with Subio platform and R gplots package. Primers used in this study were listed in Supplementary Table 2.

### Cross section

Hypocotyls of 10- or 11-day-old seedlings were fixed with FAA (formalin:acetic acid:alcohol, 1:1:18) for 1 day. Fixed hypocotyls were subjected to an ethanol series (50%, 70%, 80%, 90%, 99.5%) each for 30 minutes and then transferred into Technovit 7100 solution without Hardener II for 1 day. After the pre-incubation, samples were embedded in a mixture of Technovit 7100 + Hardener II (12.5:1) and incubated at 37 °C for more than 1 hour to harden. Technovit samples were sliced into 2 μm sections using a LEICA RM2255 microtome and stained with 0.1% toluidine blue to enable CCs to be distinguished from SEs under microscopy. Cross sections for GUS-stained samples were made as reported previously^*24*^.

### VISUAL-CC

Please see Supplementary Method

## Acknowledgements

We thank Yuki Fukaya, Akiho Suizu and Yukiko Sugisawa for technical assistance, and Enrico Scarpella, Thomas Greb, Ho-Ming Chen and Hao Yu for sharing materials.

## Author contributions

Y.K. designed the experiment. T.T., S.O., M.N., Y.O., F. T., M.S., M.S., M.W., K.T. and Y.K. performed the experiments. H.F. and Y.H shared materials and information. T.T., H.F and Y.K. wrote the manuscript.

## Data and material availability

Accession number of microarray data is GSE141037. The supplementary materials contain additional data.

---

**Supplementary Fig. 1| *smxl4 smxl5* mutants suppressed CC differentiation in VISUAL-CC**

**a**, Schematic of the VISUAL differentiation process in the WT and *smxl4 smxl5*. The *smxl4 smxl5* double mutants were known to inhibit phloem differentiation in VISUAL. **b**, SUC2 expression at 4d after VISUAL-CC induction in the WT and *smxl4 smxl5*. Asterisks indicate significant differences using the Student’s t-test (\**P* < 0.05, n = 3).

**Supplementary Fig. 2| Raw data from time-course analysis of *pSUC2:ELUC* plants**

An example of *pSUC2:ELUC* signals from individual samples is shown. Vertical axis indicates photon counts per second detected by the luminometer. Samples were classified based on LUC intensity.

**Supplementary Fig. 3| Characterization and molecular function of VC genes**

**a**, Schematic of the selection process used to identify VISUAL-CC inducible genes. Expression levels of vascular-specific genes were determined using VISUAL-CC microarray data. **b**, Expression patterns of VC genes in the root stele obtained from a transcriptome dataset^14^. Mean values from Fig. 2A are shown. **c**, Functional classification of VC genes and VPP genes. Enrichment scores were calculated using David (https://david.ncifcrf.gov/). Transporter-related genes are over-represented in this category.

**Supplementary Fig. 4 | Statistical differences in expression levels of CC-related genes among the S, M, and V samples**

Expression levels of *CEPR, FTIP1, MYR1, NaKR1,* and *SULTR2:1* were quantified using qRT-PCR and compared statistically among the S3-5, M3-5, and V3-5 samples. Relative expression levels were calculated when the expression in S3 was set to 1. Statistical differences between samples are indicated by different letters (ANOVA, Tukey-Kramer method; n = 3; error bars indicate SD).

**Supplementary Fig. 5 | Correlation analysis in microarray data between the S and M samples**

**a**, VC genes (67) vs VX genes (137) **b**, VC genes (67) vs VS genes (218) **c**, VC genes (67) vs bikinin-suppressed genes (113). The Pearson correlation coefficient and *P*-value are marked above the chart. Error bars indicate SD.

**Supplementary Fig. 6| Statistical differences in expression levels between the S and M samples**

**a**, Expression levels of *SUC2* (as CC), *SAPL* (also as CC), *SEOR1* (as SE), *APL* (as CC+SE), and *IRX3* (XY) were quantified using qRT-PCR and compared statistically between the S and M samples. Asterisks indicate significant differences using the Student’s t-test (\*\**P* < 0.005; \**P* < 0.05). **b**, Expression levels of GSK3 activity-dependent genes were quantified using qRT-PCR and compared statistically in the S and M samples. Asterisks indicate significant differences determined using the Student’s t-test (\*\**P* < 0.005; \**P* < 0.05).

**Supplementary Fig. 7| Auxin has only marginal effects on the formation of the SE-CC complex**

**a**, Time-course of *pSUC2:ELUC* signal intensities in VISUAL-CC cultures containing different concentrations of auxin (mg/L). **b** and **c**, Expression levels of *SUC2* (b) and *SEOR1* (c) in VISUAL-CC samples from cultures containing different concentrations of auxin. There are no significant differences (ANOVA, Tukey-Kramer method; n = 6; error bars indicate SD).

**Supplementary Fig. 8| Heat map of expression levels of BR biosynthesis-related genes in S and M samples.**

The upper panel shows a heat map of expression levels of 6 BR biosynthesis-related genes, which are downregulated by bikinin, in S and M samples. The lower panel indicates the mean value for each sample.

**Supplementary Fig. 9| Distribution of the SE/CC ratio among samples**

Box plots of Fig. 4e and Fig. 5c were shown. Median values were indicated by central lines. First (Q1) and third (Q3) quartile were shown as a box. Lines show the range of Q1+1.5x interquartile and Q3-1.5x interquartile. Dots indicated distributions of each plot.

**Supplementary Fig. 10| Expression pattern of GSK3s in VISUAL and VISUAL-CC transcriptome data**

**a**, Normalized expression levels of procambium (*AtHB8*), xylem *(IRX3*), phloem SE (*CALS7*) and SKI/II GSK3 subgroup genes in VISUAL transcriptome data. Error bars indicate SD (n=3). **b,** Normalized expression levels of procambium (*AtHB8*), xylem *(IRX3*), phloem SE (*CALS7*) and SKI/II GSK3 subgroup genes in VISUAL-CC transcriptome data.

**Supplementary Fig. 11 | Effect of brassinolide treatment on phloem development**

**a**, Toluidine blue-stained transverse sections of mock-treated (DMSO) and bikinin-treated hypocotyls. SE: white empty cell; CC: dense purple cell. **b**, SE/CC ratios (%) in the WT treated with none (control), 100 nM BL, and 1000 nM BL were calculated from toluidine blue-stained sections (n = 11–14). Numbers of individuals are marked. **c**, Box plot of (b) was shown. Scale bars: 10 μm.

**Supplementary Fig. 12 | Transverse sections of *bes1 bzr1* mutants**

Toluidine blue-stained transverse sections for 11-day-old hypocotyls of WT, *bes1-D bzr1-D* (gain-of-function), and *bes1-1 bzr1-2* (loss-of-function) mutant plants. Scale bars: 50 μm.

