## Supplementary information for "GSK3 activity is a cell fate switch that balances the ratio of vascular cell type"

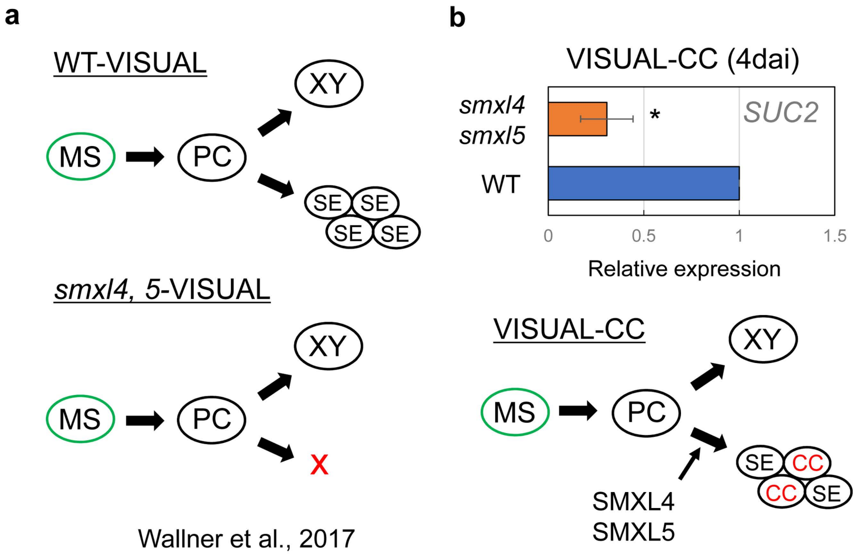


**Supplementary Fig. 1| *smxl4 smxl5* mutants suppressed CC differentiation in VISUAL-CC**
**a**, Schematic of the VISUAL differentiation process in the WT and *smxl4 smxl5*. The *smxl4 smxl5* double mutants were known to inhibit phloem differentiation in VISUAL. **b**, SUC2 expression at 4d after VISUAL-CC induction in the WT and *smxl4 smxl5*. Asterisks indicate significant differences using the Student's t-test (**P* < 0.05, n = 3).

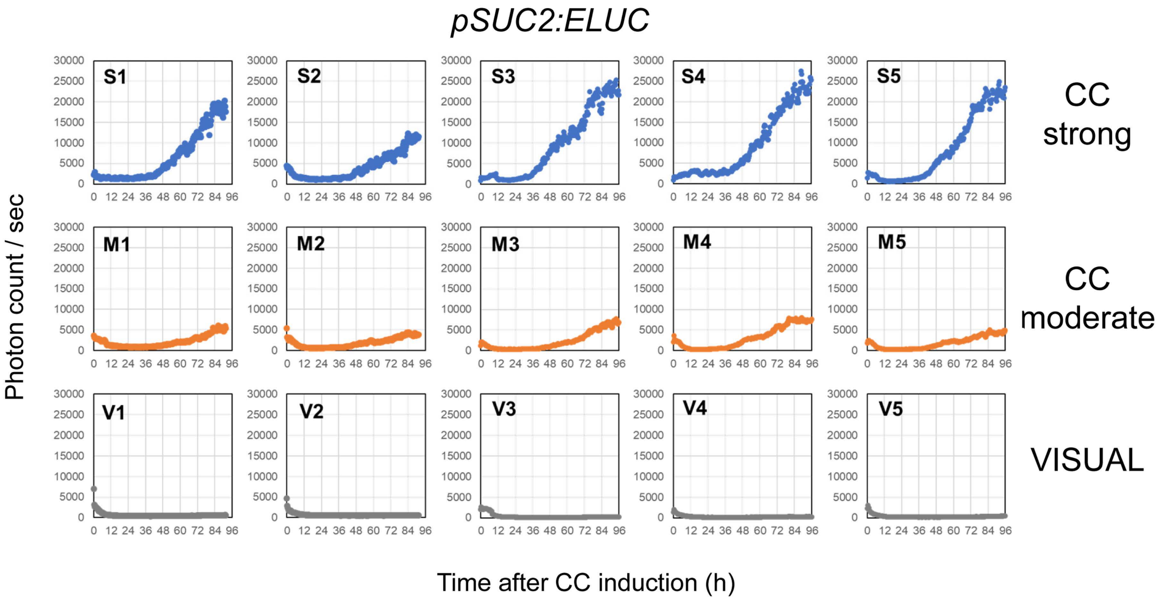


**Supplementary Fig. 2| Raw data from time-course analysis of *pSUC2:ELUC* plants**
An example of *pSUC2:ELUC* signals from individual samples is shown. Vertical axis indicates photon counts per second detected by the luminometer. Samples were classified based on LUC intensity.

**a**

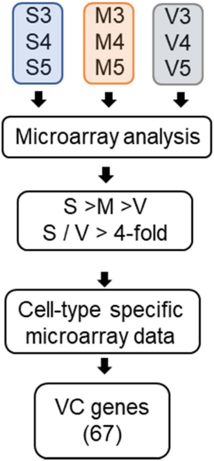


**b**

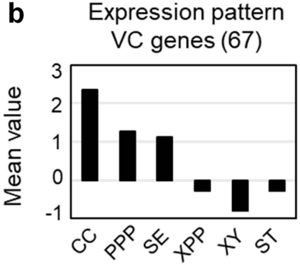


**c**

Gene functional classification

| Gene functional clustering | Enrichment score |
|---|---|
| Peroxidase (4) | 2.12 |
| **Transporter (10)** | **0.82** |
| Transcription factor(8) | 0.57 |

**Supplementary Fig. 3| Characterization and molecular function of VC genes**
**a**, Schematic of the selection process used to identify VISUAL-CC inducible genes. Expression levels of vascular-specific genes were determined using VISUAL-CC microarray data. **b**, Expression patterns of VC genes in the root stele obtained from a transcriptome dataset[14]. Mean values from Fig. 2A are shown. **c**, Functional classification of VC genes and VPP genes. Enrichment scores were calculated using David (https://david.ncifcrf.gov/). Transporter-related genes are over-represented in this category.

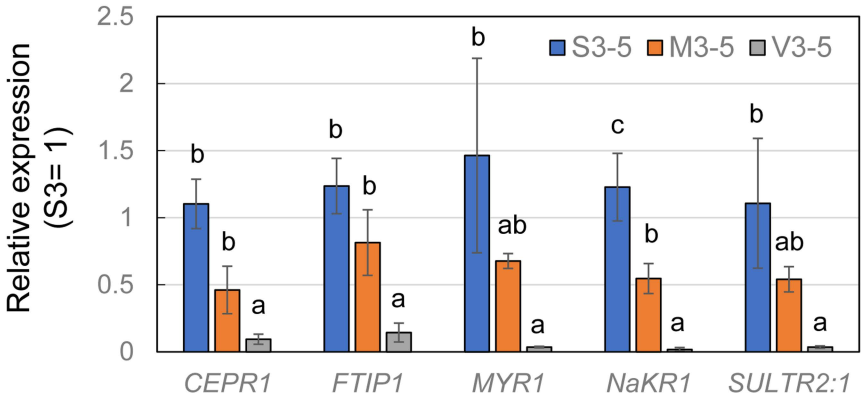


**Supplementary Fig. 4 | Statistical differences in expression levels of CC-related genes among the S, M, and V samples**

Expression levels of *CEPR, FTIP1, MYR1, NaKR1,* and *SULTR2:1* were quantified using qRT-PCR and compared statistically among the S3-5, M3-5, and V3-5 samples. Relative expression levels were calculated when the expression in S3 was set to 1. Statistical differences between samples are indicated by different letters (ANOVA, Tukey-Kramer method; n = 3; error bars indicate SD).

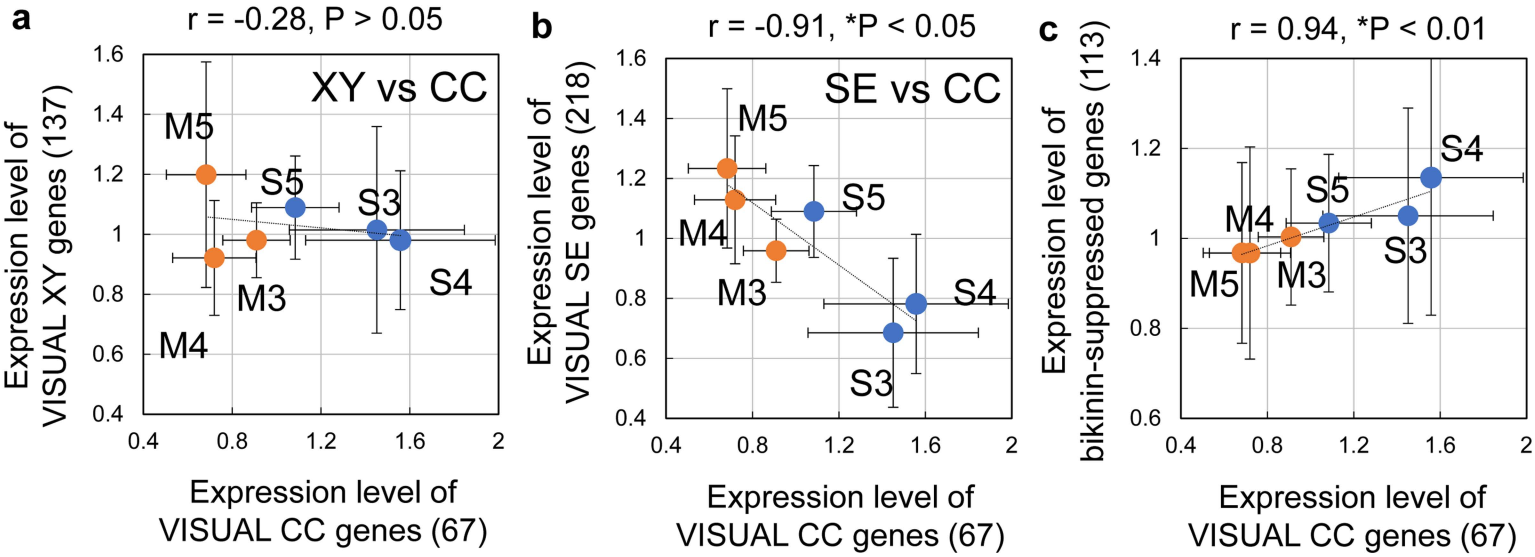


**Supplementary Fig. 5 | Correlation analysis in microarray data between the S and M samples**

**a**, VC genes (67) vs VX genes (137) **b**, VC genes (67) vs VS genes (218) **c**, VC genes (67) vs bikinin-suppressed genes (113). The Pearson correlation coefficient and *P*-value are marked above the chart. Error bars indicate SD.

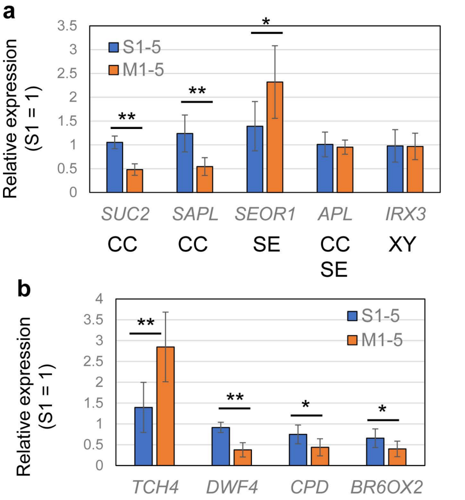


**Supplementary Fig. 6| Statistical differences in expression levels between the S and M samples**

**a**, Expression levels of *SUC2* (as CC), *SAPL* (also as CC), *SEOR1* (as SE), *APL* (as CC+SE), and *IRX3* (XY) were quantified using qRT-PCR and compared statistically between the S and M samples. Asterisks indicate significant differences using the Student's t-test (**$P$ < 0.005; *$P$ < 0.05). **b**, Expression levels of GSK3 activity-dependent genes were quantified using qRT-PCR and compared statistically in the S and M samples. Asterisks indicate significant differences determined using the Student's t-test (**$P$ < 0.005; *$P$ < 0.05).

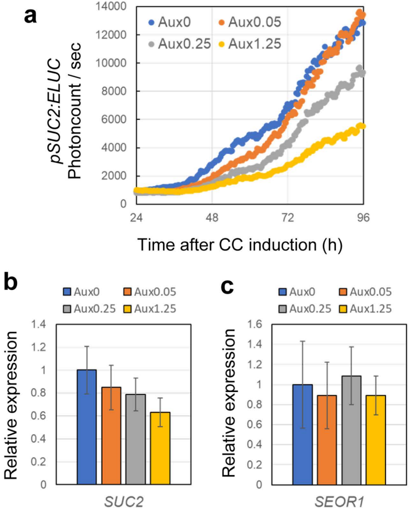


**Supplementary Fig. 7| Auxin has only marginal effects on the formation of the SE-CC complex**

**a**, Time-course of *pSUC2:ELUC* signal intensities in VISUAL-CC cultures containing different concentrations of auxin (mg/L). **b** and **c**, Expression levels of *SUC2* (b) and *SEOR1* (c) in VISUAL-CC samples from cultures containing different concentrations of auxin. There are no significant differences (ANOVA, Tukey-Kramer method; n = 6; error bars indicate SD).

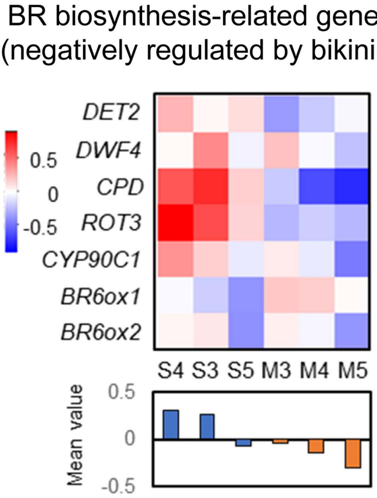


**Supplementary Fig. 8| Heat map of expression levels of BR biosynthesis-related genes in S and M samples.**

The upper panel shows a heat map of expression levels of 6 BR biosynthesis-related genes, which are downregulated by bikinin, in S and M samples. The lower panel indicates the mean value for each sample.

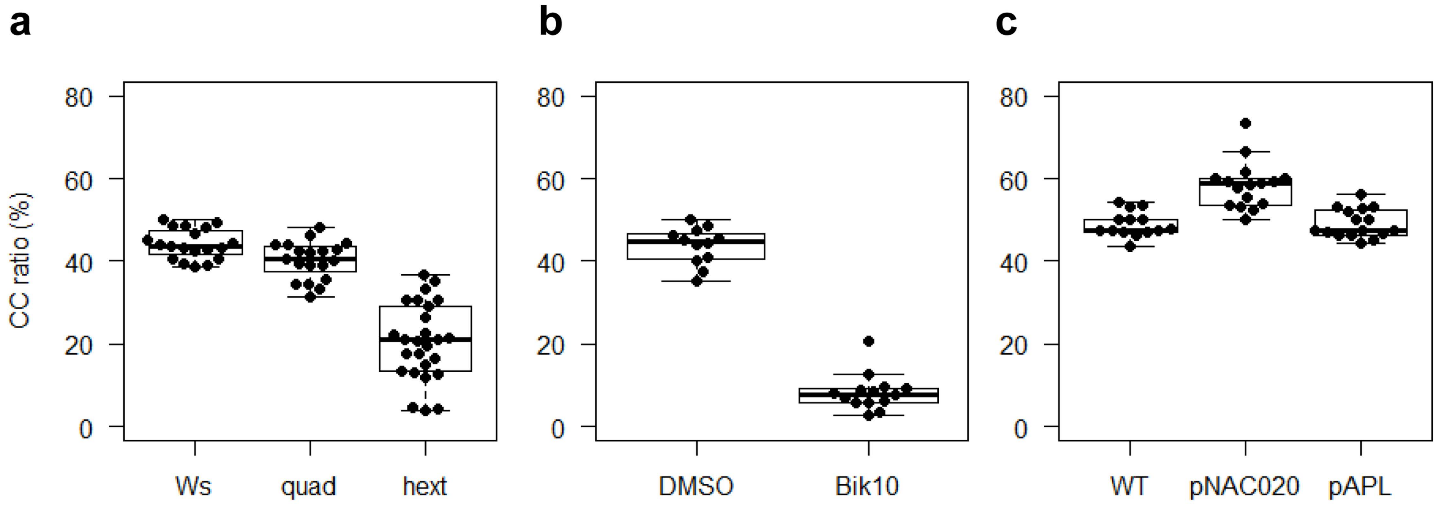


**Supplementary Fig. 9| Distribution of the SE/CC ratio among samples**
Box plots of Fig. 4e and Fig. 5c were shown. Median values were indicated by central lines. First (Q1) and third (Q3) quartile were shown as a box. Lines show the range of Q1+1.5x interquartile and Q3-1.5x interquartile. Dots indicated distributions of each plot.

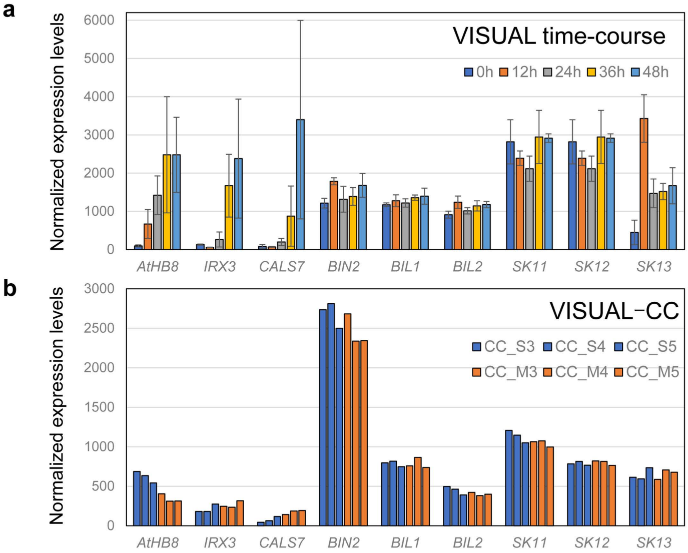


**Supplementary Fig. 10| Expression pattern of GSK3s in VISUAL and VISUAL-CC transcriptome data**

**a**, Normalized expression levels of procambium (*AtHB8*), xylem (*IRX3*), phloem SE (*CALS7*) and SKI/II GSK3 subgroup genes in VISUAL transcriptome data. Error bars indicate SD (n=3). **b,** Normalized expression levels of procambium (*AtHB8*), xylem (*IRX3*), phloem SE (*CALS7*) and SKI/II GSK3 subgroup genes in VISUAL-CC transcriptome data.

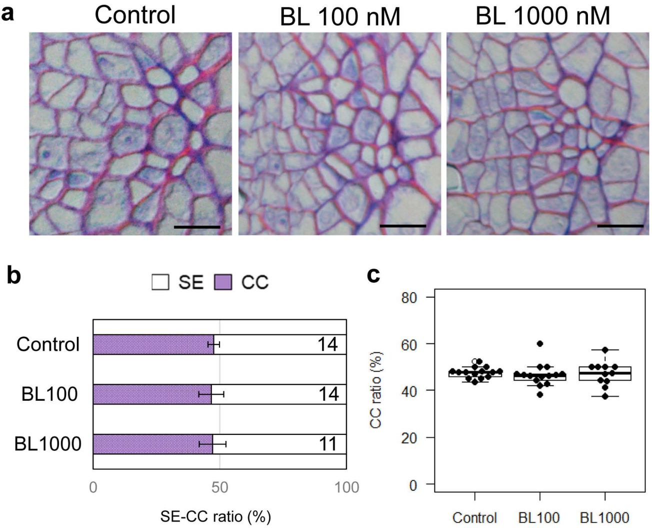


**Supplementary Fig. 11 | Effect of brassinolide treatment on phloem development**

**a**, Toluidine blue-stained transverse sections of mock-treated (DMSO) and bikinin-treated hypocotyls. SE: white empty cell; CC: dense purple cell. **b**, SE/CC ratios (%) in the WT treated with none (control), 100 nM BL, and 1000 nM BL were calculated from toluidine blue-stained sections (n = 11– 14). Numbers of individuals are marked. **c**, Box plot of (b) was shown. Scale bars: 10 μm.

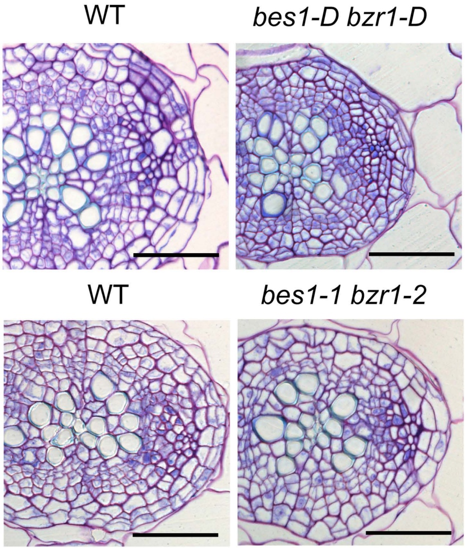


**Supplementary Fig. 12 | Transverse sections of *bes1 bzr1* mutants**

Toluidine blue-stained transverse sections for 11-day-old hypocotyls of WT, *bes1-D bzr1-D* (gain-of-function), and *bes1-1 bzr1-2* (loss-of-function) mutant plants. Scale bars: 50 μm.

Supplementary Table 1. The list of VC genes (67).

| AGI code | Description (based on TAIR) |
|---|---|
| At1g01470 | Late embryogenesis abundant protein (LEA14) |
| At1g10380 | Putative membrane lipoprotein |
| At1g12090 | extensin-like protein (ELP) |
| At1g13380 | Protein of unknown function (DUF1218) (DUF1218) |
| At1g13590 | phytosulfokine 1 precursor (PSK1) |
| At1g22710 | sucrose-proton symporter 2 (SUC2) |
| At1g49310 | transmembrane protein |
| At1g49500 | transcription initiation factor TFIID subunit 1b-like protein |
| At1g59740 | NRT1/ PTR FAMILY 4.3 |
| At1g59960 | NAD(P)-linked oxidoreductase superfamily protein |
| At1g68740 | PHO1;H1 |
| At1g76130 | alpha-amylase-like 2 (AMY2) |
| At1g77380 | amino acid permease 3 (AAP3) |
| At2g02020 | NRT1/ PTR FAMILY 8.4 |
| At2g02130 | low-molecular-weight cysteine-rich 68 (LCR68) |
| At2g04160 | Subtilisin-like serine endopeptidase family protein (AIR3) |
| At2g19590 | ACC oxidase 1 (ACO1) |
| At2g22860 | phytosulfokine 2 precursor (PSK2) |
| At2g30070 | potassium transporter 1 (KT1) |
| At2g37130 | Peroxidase superfamily protein |
| At2g44380 | Cysteine/Histidine-rich C1 domain family protein |
| At2g46690 | SMALL AUXIN UPREGULATED RNA 32 (SAUR32) |
| At3g09260 | BGLU23 |
| At3g12730 | SAPL |
| At3g12750 | zinc transporter 1 precursor (ZIP1) |
| At3g14560 | hypothetical protein |
| At3g14840 | LYSM RLK1-INTERACTING KINASE 1 (LIK1) |
| At3g15950 | DNA topoisomerase-related |
| At3g16450 | JACALIN-RELATED LECTIN 33 (JAL33) |
| At3g16460 | JACALIN-RELATED LECTIN 34 (JAL34) |
| At3g20370 | TRAF-like family protein |
| At3g21770 | Peroxidase superfamily protein |
| At3g23050 | indole-3-acetic acid 7 (IAA7) |
| At3g60720 | plasmodesmata-located protein 8 (PDLP8) |
| At3g63110 | isopentenyltransferase 3 (IPT3) |
| At4g12470 | azelaic acid induced 1 (AZI1) |
| At4g12550 | Auxin-Induced in Root cultures 1 (AIR1) |
| At4g14465 | AT-hook motif nuclear-localized protein 20 (AHL20) |
| At4g15660 | GRXS8 |

| | |
|---|---|
| At4g15690 | GRXS5 |
| At4g19840 | phloem protein 2-A1 (PP2-A1) |
| At4g21960 | Peroxidase superfamily protein |
| At4g27410 | NAC (No Apical Meristem) domain transcriptional regulator superfamily protein (RD26) |
| At4g32290 | Core-2/I-branching beta-1,6-N-acetylglucosaminyltransferase family protein |
| At4g32870 | Polyketide cyclase/dehydrase and lipid transport superfamily protein |
| At4g35480 | RING-H2 finger A3B (RHA3B) |
| At4g36410 | ubiquitin-conjugating enzyme 17 (UBC17) |
| At4g37540 | LOB domain-containing protein 39 (LBD39) |
| At5g01210 | HXXXD-type acyl-transferase family protein |
| At5g01840 | ovate family protein 1 (OFP1) |
| At5g02260 | expansin A9 (EXPA9) |
| At5g02600 | NaKR1 |
| At5g07010 | sulfotransferase 2A (ST2A) |
| At5g18240 | myb-related protein 1 (MYR1) |
| At5g23820 | MD2-RELATED LIPID RECOGNITION 3 (ML3) |
| At5g24800 | BASIC LEUCINE ZIPPER 9 (BZIP9) |
| At5g26260 | TRAF-like family protein |
| At5g26280 | TRAF-like family protein |
| At5g28770 | BASIC LEUCINE ZIPPER 63 (BZIP63) |
| At5g43380 | type one serine/threonine protein phosphatase 6 (TOPP6) |
| At5g43580 | UNUSUAL SERINE PROTEASE INHIBITOR (UPI) |
| At5g49660 | CEPR1 / XIP1 |
| At5g54130 | Calcium-binding endonuclease/exonuclease/phosphatase family |
| At5g59080 | hypothetical protein |
| At5g63710 | Leucine-rich repeat protein kinase family protein |
| At5g64120 | Peroxidase superfamily protein |
| At5g65970 | Seven transmembrane MLO family protein (MLO10) |

**Supplementary Table 2**

Primers used in this study for qRT-PCR

| name | sequence (5'-3') |
|---|---|
| CPD-L | AACCCTTGGAGATGGCAGA |
| CPD-R | GTAACCGGGACATAGCCTTG |
| DWF4-L | TTCTCGTTATGACCAACCTAATCTC |
| DWF4-R | AGGATGACGCTCCGTTGTT |
| UBQ14-L | TCCGGATCAGCAGAGGTT |
| UBQ14-R | TCTGGATGTTGTAGTCAGCAAGA |
| APL-L | TGGATATTCAGCGCAACGTA |
| APL-R | TGCACTTCCATTTGCATCTC |
| SUC2-L | TAGCCATTGTCGTCCCTCA |
| SUC2-R | CCACCACCGAATAGTTCGTC |
| IRX3-L | TGACATGAATGGTGACGTAGC |
| IRX3-R | CATCAAATGCTCCTTATCACCTT |
| SEOR1-L | AAGACACCAACGCCTCCA |
| SEOR1-R | CGATAGCATAGGAGACACTATCAAGA |
| CALS7-L | GCAGTAATGGAACTCCCTGAGA |
| CALS7-R | GGCTGAATGGAATCTTGGTC |
| SAPL-L | AGAGCCATCTCCAGAAGTTCA |
| SAPL-R | CCTTCGAAGATCCAACATGG |
| TCH4-L | GCTCAACAAAGGATGAGATGG |
| TCH4-R | CCTCTTCGCATCCGTACAAT |
| BR6ox2-L | CCCATGGAGATGGATGGA |
| BR6ox2-R | CTTTCCAGGGCAAAGCCTA |
| SULTR2:1-L | AACGATCTCATGGCTGGTTTA |
| SULTR2:1-R | TTGCATAACCAATGCTCTGC |
| NaKR1-L | GCTCAGTTTTGGCCTGAGATT |
| NaKR1-R | GTGGTGAATCAGCCAGTCCT |
| CEPR1-L | TATGGCTGGCACCTATGGTT |
| CEPR1R | GATCGTTGCTTTGGACGAGT |
| FTIP1-L | GCGCAAGATGTTGAGCCTA |
| FTIP1-R | TTGTACTTTAACGAAAGCTTGAGG |
| MYR1-L | GAAGTAGACGAAAGTCACAGTGAGAG |
| MYR1-R | GGCATCACTTATGGGTAAGTTCA |

Supplementary Method

**VISUAL-CC**

VISUAL-CC consists of two distinct steps; vascular stem cell formation and subsequent phloem differentiation. As the initial step, 6 or 7 day-old seedlings were cultured with the conventional VISUAL medium for 2 days in order to induce sufficient amount of (pro)cambial cells. After that samples are transferred into VISUAL-CC medium for SE-CC complex differentiation.

<u>**Materials**</u>

**Growth of plant samples before VISUAL induction**

1. MS growth medium: It contains 2.2 g/L MS Basal Medium (Sigma), 10 g/L sucrose and 0.5 g/L 2-morpholinoethanesulfonic acid monohydrate (MES) in Milli-Q water and the pH is adjusted to 5.7 with KOH. The solution is autoclaved at 120°C for 20 min and can be stored at room temperature up to several weeks.
2. Sterilizing solution: Sodium hypochlorite solution is diluted in Milli-Q water in the ratio 1:9 (v/v) and 0.1% of Triton-X100 is added. This solution is prepared immediately before the sterilizing procedure.
3. Sterilized 6-well plate (Sumilon)
4. Autoclaved Milli-Q water
5. Surgical tape
6. Continuous light chamber (22°C, 45–55 μmol/$m^2$/s)
7. Rotary shaker (Taitec)

**VISUAL and VISUAL-CC**

1. VISUAL base medium: It contains 2.2 g/L MS Basal Medium and 50 g/L D(+)-Glucose in Milli-Q water and the pH is adjusted to 5.7 with KOH The solution is autoclaved at 120°C for 20 min and can be stored at room temperature for several weeks.
2. VISUAL-CC base medium: It contains 2.2 g/L MS Basal Medium and 10 g/L D(+)-Glucose in Milli-Q water and the pH is adjusted to 5.7 with KOH The solution is autoclaved at 120°C for 20 min and can be stored at room temperature for several

weeks. Note that Glucose concentration is different from that of VISUAL base medium.

3. 2,4-D stock: 2.5 g/L 2,4-D stock dissolved in autoclaved Milli-Q water and sterilized through 0.22 µm filter units. Stored in small amounts in sampling tubes at -20°C.
4. Kinetin stock: 0.5 g/L Kinetin stock dissolved in 0.1 M KOH and sterilized through 0.22 µm filter units. Stored in small amounts in sampling tubes at -20°C.
5. Bikinin stock: 10 mM Bikinin stock dissolved in DMSO and sterilized through 0.22 µm filter units. Stored in small amounts in sampling tubes at -20°C.
6. Sterilized 12-well plate (Sumilon)
7. Surgical forceps
8. Continuous light chamber (22°C, 60–70 µmol/m$^2$/s)
9. Rotary shaker (Taitec)

### Methods

#### Growth of plant samples before VISUAL induction

1. Sterilizing solution is added to the Arabidopsis seeds in 1.5 mL sampling tubes and gently mixed using a rotator for 5 min. The tubes are transferred inside a clean bench and allowed to stand for further 5 min. The sterilizing solution is then removed using a pipette and the seeds are washed with autoclaved Milli-Q water three times. The seeds are soaked in water at 4°C for 2 days to keep the germination timing constant.
2. 10 mL of the prepared MS growth medium is poured into each well of a 6-well plate. Seeds are sown at a density of 8-10 seeds/well containing the MS growth medium and the plate is sealed with surgical tape. The well plate is incubated for 6-7 days under continuous light (22°C, 45–55 µmol/m$^2$/s) with shaking at 110 rpm on a rotary shaker.

#### VISUAL

1. 2,4-D stock, kinetin stock and bikinin stock are defrosted at room temperature before use. The tubes are transferred inside a clean bench and added to the VISUAL base medium to obtain a final concentration of 1.25 mg/L 2,4-D, 0.25 mg/L kinetin and 10 µM bikinin. About 2.5 mL of the above medium is then added into each well of a

12-well plate.

2. A pair of sharp surgical forceps are used to cut the bottom half of Arabidopsis 6-7 day-old plants across the center of the hypocotyl and the roots are removed. About 4 of the Arabidopsis explants are then transferred carefully to each well containing the induction medium using forceps and the 12-well plate sealed with surgical tape. The explants are cultured for 2 days under continuous light (22°C, 60–70 μmol/$m^2$/s) with shaking at 110 rpm on a rotary shaker.

**VISUAL-CC**

1. 2,4-D stock, kinetin stock and bikinin stock are defrosted at room temperature before use. The tubes are transferred inside a clean bench and added to the VISUAL-CC base medium to obtain a final concentration of 0.25 mg/L 2,4-D, 0.25 mg/L kinetin and 1 μM bikinin. About 2.5 mL of the above medium is then added into each well of a 12-well plate. Note that auxin and bikinin concentration is decreased when compared to the VISUAL.
2. VISUAL-induced samples were transferred into the new CC medium and then cultured for 4 days under dark condition (22°C) with shaking at 110 rpm on a rotary shaker. Note that light severely affect CC differentiation ratio.
